# Beta-Band Oscillations without Segregated Pathways: the opposing Roles of D2 and D5 Receptors in the Basal Ganglia

**DOI:** 10.1101/161661

**Authors:** Jean F. Liénard, Lise Aubin, Ignasi Cos, Benoît Girard

## Abstract

The timescales of the dynamics of a system depend on the combination of the timescales of its components and of its transmission delays. Parkinson’s disease is characterized by the death of dopaminergic neurons and the emergence of strong *β*-band (15-35Hz) oscillations throughout the basal ganglia nuclei. Here we combine experimental stimulation data from ten studies, that reveal the timing of excitatory and inhibitory events in the basal ganglia circuit, to estimate its set of transmission delays. In doing so, we reveal possible inconsistencies in the existing data, calling for replications, and we propose two possible sets of transmission delays.

We then integrate these delays in a model of the primate basal ganglia, that does not rely on direct and indirect pathways’ segregation, and show that, while much attention has been given to the role of the striatal dopaminergic receptors in Parkinson’s disease symptoms, extrastriatal dopaminergic depletion in the external part of the globus pallidus and in the subthalamic nucleus is sufficient to generate *β*-band oscillations in the high part of the band. More specifically, we show that that D2 and D5 dopamine receptors in these nuclei play opposing roles in the emergence of *β*-band oscillations, thereby explaining how completely deactivating D5 receptors in the subthalamic nucleus can, paradoxically, cancel oscillations.

## 1. Introduction

Along with bradykinesia, hypokinesia, akinesia and resting tremor, one of the major hallmarks of Parkinson’s Disease (PD) is the aberrant *β*-band (15-35 Hz) oscillatory activity recorded in several nuclei of the Basal Ganglia (Marsden, 1984, 1989, Berardelli et al., 1996, Samii et al., 2004, Berardelli et al., 2001, Mazzoni et al., 2012). Specifically, stronger than normal *β*-band power has consistently been revealed in EEG and MEG recordings from PD patients and in primate models of PD (Oswal et al., 2013), and is also found in electro-physiological recordings in the sub-thalamic nucleus (STN) and globus pallidus (GP) (Filion and Tremblay, 1991, Nini et al., 1995, Levy et al., 2000, Brown et al., 2001, Kühn et al., 2006, Weinberger et al., 2009). Remarkably, the fact that *β*-band activity can be restored to normal levels by administration of the DA precursor L-dopa (Brown and Marsden, 1999, Doyle et al., 2005), shows that DA and the intensity of neural activity in the *β*-band are intimately related (Oswal et al., 2013). However, the specifics of their relationship remain elusive. Traditionally, a change in the balance between the direct and indirect pathways of the basal ganglia (BG) (Albin et al., 1989, Gurney et al., 2001, Frank et al., 2004, Frank, 2005) has been considered to be the origin of the anti-kinetic PD symptoms, later extending to encompass the generation of abnormal oscillations (Humphries et al., 2006, Van Albada and Robinson, 2009, Van Albada et al., 2009, Tsirogiannis et al., 2010, Kumar et al., 2011, Lindahl and Hellgren Kotaleski, 2016).

Although frequent in the computational literature, there is growing experimental evidence suggesting this hypothesis might need partial revision (referred to as the second problem on the basal ganglia, in Nambu, 2008). Specifically, in order to cause this imbalance, striatal medium spiny neurons (MSN) would have to be organized into independent populations that express either D1 or D2 receptors, each then respectively projecting to the internal or external segments of the globus pallidus (GPi/GPe). While striatal pathways may indeed be segregated in mice (e.g., Valjent et al. 2009, but see also Cazorla et al. 2014), tracing studies in monkeys (Parent et al., 1995, Lévesque and Parent, 2005) and rats (Kawaguchi et al., 1990, Wu et al., 2000, Fujiyama et al., 2011) have consistently shown that the majority of MSN projects both to the GPe and GPi.

Thus, at least for monkeys, and possibly for rats, there is considerable anatomical evidence that casts a doubt on the position that these pathways act and interact independently, and that consequently their functional imbalance is the cause of *β*-band oscillations in the BG. Alternatively, stronger than normal *β*-band oscillations have also been linked to an imbalance of extra-striate DA receptors (Benazzouz et al., 2014), which are present across all basal ganglia nuclei (Rommelfanger and Wichmann, 2010) and are equally affected by DA loss. However, their roles are still not fully understood.

In this study we test the hypothesis that extra-striate DA receptors are sufficient to set the basal ganglia in an oscillatory regime under dopamine depletion, by means of a computational model. To do so we use an existing BG model of the macaque monkey (Liénard and Girard, 2014, see Fig. 1) that we extend with transmission delays as realistic as possible, so as to properly estimate the frequency of these oscillations. The estimation of these delays is based on ten studies (Yoshida et al., 1993, Nambu et al., 2000, Turner and DeLong, 2000, Nambu et al., 2002, Kita et al., 2004, 2006, Tachibana et al., 2008, Iwamuro et al., 2009, 2017, Polyakova et al., 2020) that provide the latency of the excitatory and inhibitory events in all the BG following stimulations applied in the cortex, the striatum, the subthalamic nucleus or the external globus pallidus. The combination of all these data reveal some possibly contradictory measurements, leading us to use two sets of delays. The first one is the best possible compromise that can be established when trying to satisfy all the –possibly contradictory– constraints, while the second one is obtained by removing the data from a single study, so as to minimize experimental data incompatibilities.

**Figure 1:**
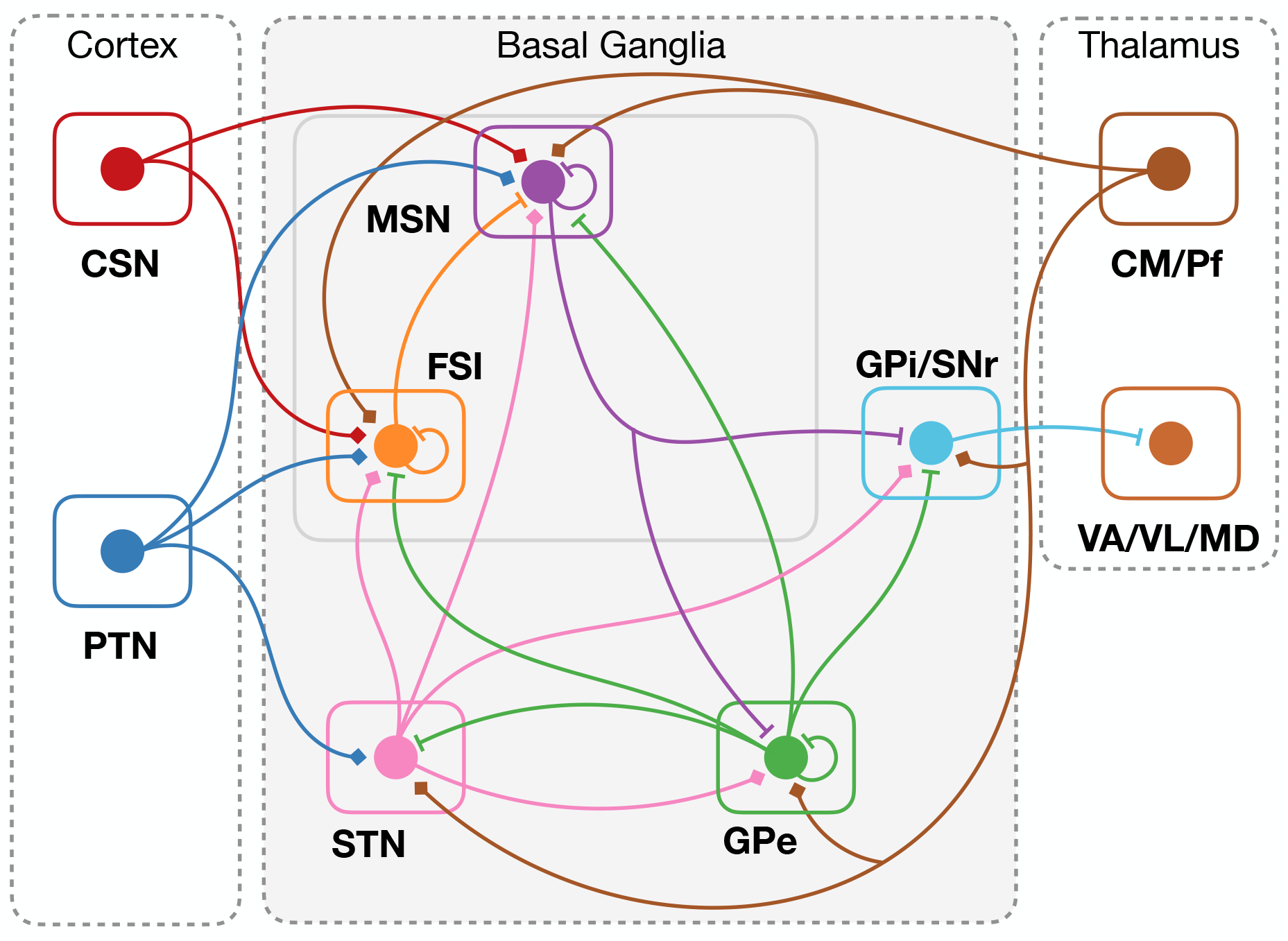
The simulated populations of the Liénard and Girard (2014) basal ganglia model (gray back-ground) and their interconnections. The cortical and thalamic populations (white backgroud) are not explicitly simulated: the cortico-striatal neurons (CSN), the pyramidal tract neurons (PTN) and the centro-median/parafascicular neurons (CM/Pf) are external inputs; the ventro-anterior, ventro-lateral and medio-dorsal neurons of the thalamus (VA/VL/MD) are the targets of the GPi/SNr output of the model. Diamond projections are excitatory, flat projections are inhibitory. All figures by Aubin, Liénard & Girard (2022); available under a CC-BY4.0 licence (https://doi.org/10.6084/m9.figshare.21131911).

Using both sets of delays with the Liénard and Girard (2014) model, we simulated the effect of a gradual DA depletion on the dynamics of the neural circuitry by varying the only two free parameters of the model. Our main result shows that as the level of DA transits below a critical boundary, the STN-GPe network starts oscillating within the high part of the *β*-band. Specifically, our model of the BG shows that D2 dopamine receptors in the STN and the GPe, and D5 dopamine receptors in the STN, play opposing roles for the emergence of *β*-band oscillations, and are the simplest cause for this phenomenon. The robustness and simplicity of our predictions strongly support abnormal activity of extra-striate dopamine receptors as the cause of parkinsonian oscillations, and that their frequency is set by the axonal transmission delays of the STN-GPe loop.

## 2. Results

### Transmission Delays in the primate Basal Ganglia

The method we used to estimate the delays between nuclei was based on an exhaustive search, tailored to reproduce the timings recorded during stimulation experiments. All delay combinations were evaluated (taking delay values in the 1-12 ms interval), the latency of an experimentally measured inhibitory or excitatory event was simply compared to the sum of the transmission delays and neural processing time in all the possible pathways, and resulted in a score dependent on the difference between the measurement and the closest pathway prediction (for more details, see section 4.1). This operation yielded a first set of best fitting axonal delays shown in Table 1, in the column labelled *Cmpr*. The first observation deriving from the comparison of the predicted excitation and inhibition events with the data gathered from the experimental stimulation studies (fig. 2, top row), is that this set of delays predicts an unreported inhibitory event in the STN, after a cortical stimulation. This early inhibition however happens 18 ms after stimulation, only 1 ms before the expected late excitation event, and could therefore be masked by the latter and be experimentally undetectable. Indeed, inhibitory events are detected when the firing rate drops below the mean firing rate minus 1.5 standard deviation, for at least two consecutive 1 ms bins (Polyakova et al., 2020).

**Table 1:** Axonal delays in ms from previous computational studies of the primate BG and obtained with our model and fitting method. N/A (not applicable) indicates connections excluded form a given model. Cmpr: compromise solution; Antdrm: solution with antidromic Str →Ctx activations; No-Plkv: solution without the data from Polyakova et al. (2020), see text for more details

| Connection | A | B | C | D | E | F | G | H | Cmpr | Antdrm | NoPlkv |
| --- | --- | --- | --- | --- | --- | --- | --- | --- | --- | --- | --- |
| Ctx → Str | 6 | 2 | 4 | N/A | N/A | N/A | 2.5 | N/A | **6** | **9** | **9** |
| Ctx → STN | 5 | 1 | 1 | N/A | N/A | 5.5 | 2.5 | N/A | **4** | **4** | **4** |
| Str → GPe | N/A | 1 | 3 | N/A | N/A | N/A | 7 | N/A | **8** | **4** | **6** |
| Str → GPi | 10 | 1 | 3 | N/A | N/A | N/A | 7 | N/A | **11** | **8** | **8** |
| STN → GPe | N/A | 1 | 1 | 6 | 5 | 6 | 2 | 5 | **9** | **4** | **2** |
| STN → GPi | 5 | 1 | 1 | N/A | N/A | N/A | 4.5 | N/A | **4** | **4** | **4** |
| GPe → STN | N/A | 1 | 1 | 6 | 5 | 6 | 1 | 5 | **1** | **2** | **7** |
| GPe → GPi | N/A | 1 | 1 | N/A | N/A | N/A | 3 | N/A | **1** | **4** | **3/4** |

<sup>A</sup>:Leblois et al. (2006a) <sup>B</sup>:Van Albada and Robinson (2009) <sup>C</sup>:Tsirogiannis et al. (2010) <sup>D</sup>:Holgado et al. (2010) <sup>E</sup>:Kumar et al. (2011) <sup>F</sup>:Pavlidis et al. (2015) <sup>G</sup>:Lindahl and Hellgren Kotaleski (2016) <sup>H</sup>:Shouno et al. (2017)

**Figure 2:**
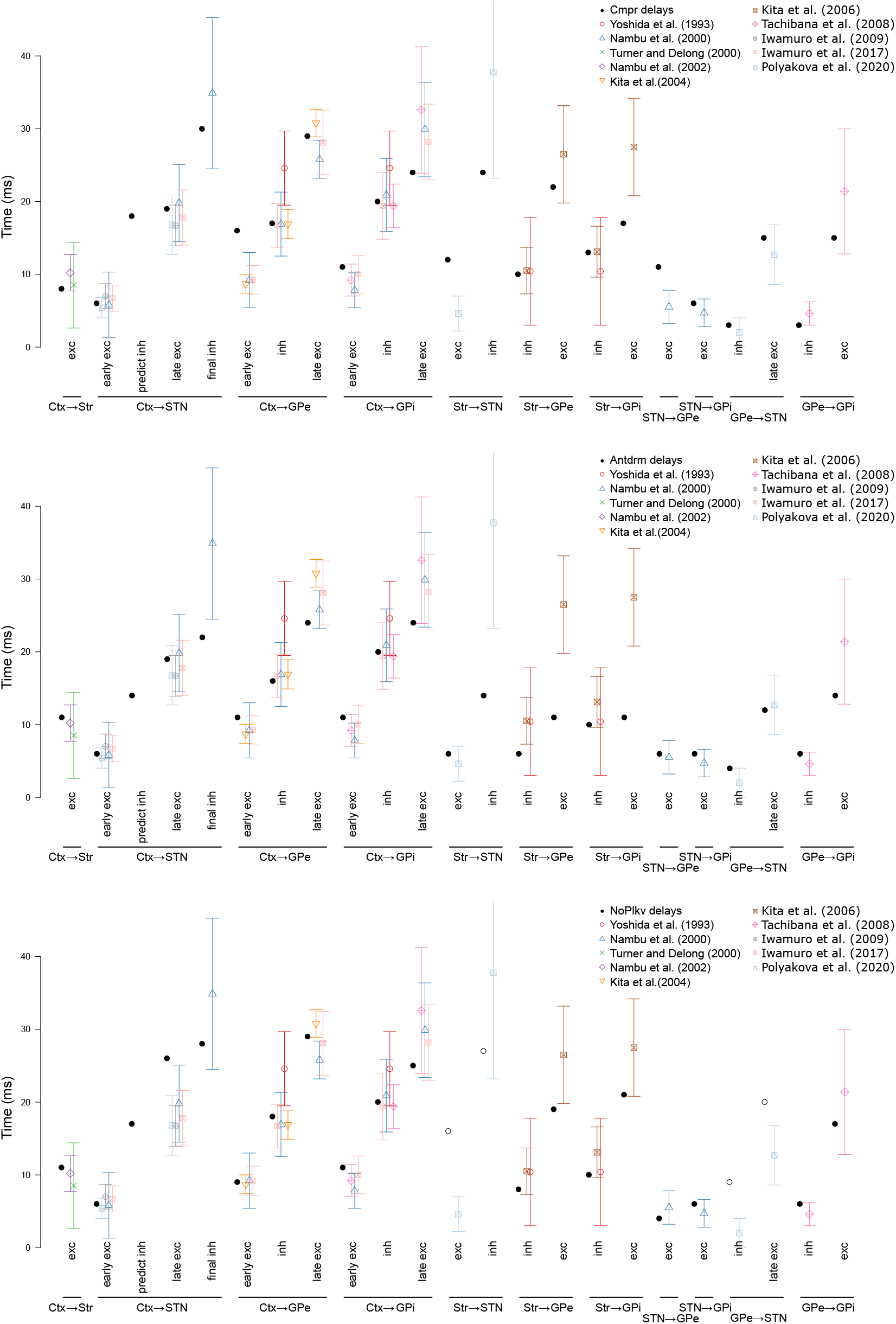
Comparison of the latencies of the excitatory and inhibitory effects measured after the stimulations (averages and standard deviations) from our set of reference studies, with those predicted by the three considered sets of delays. Top: compromise solution3b2ased on all the studies (Cmpr set); middle: solution obtained when allowing for antidromic activations of the cortex after striatum stimulation (Antdrm); bottom: solution obtained after exclusion of the (Polyakova et al., 2020) data (NoPlkv), the white circles are the latencies that were excluded from the optimization.

We can observe that the set of event timings reported in Polyakova et al. (2020) has some specificities that impose strong constraints on the results of our search (see Table 2): in this work only were measured event latencies in the STN, caused by striatal and GPe stimulations. In particular, GPe stimulations elicited an extremely fast (as it could begin immediately after the stimulation artifact) *early inhibition* effect, and striatal stimulations elicited an *early excitation*, after 4.6 ms on average. The first effect can be mediated by the direct GPe → STN connection, and indeed such a fast inhibition could not reasonably be the result of a polysynaptic pathway. This thus forces the delay of this projection to take the minimal possible value considered, 1 ms. Using such a short delay here has of course repercussions on a number the latencies of the excitatory and inhibitory events predicted by the model (fig. 2, top). Those, whose pathway use this projection, tend to be in the lower ranges: either close to the average minus one standard deviation (Ctx → GPi late excitation, Str → STN inhibition, Str → GPe excitation, GPe → GPi excitation) or even lower (Str → GPi excitation).

**Table 2:**
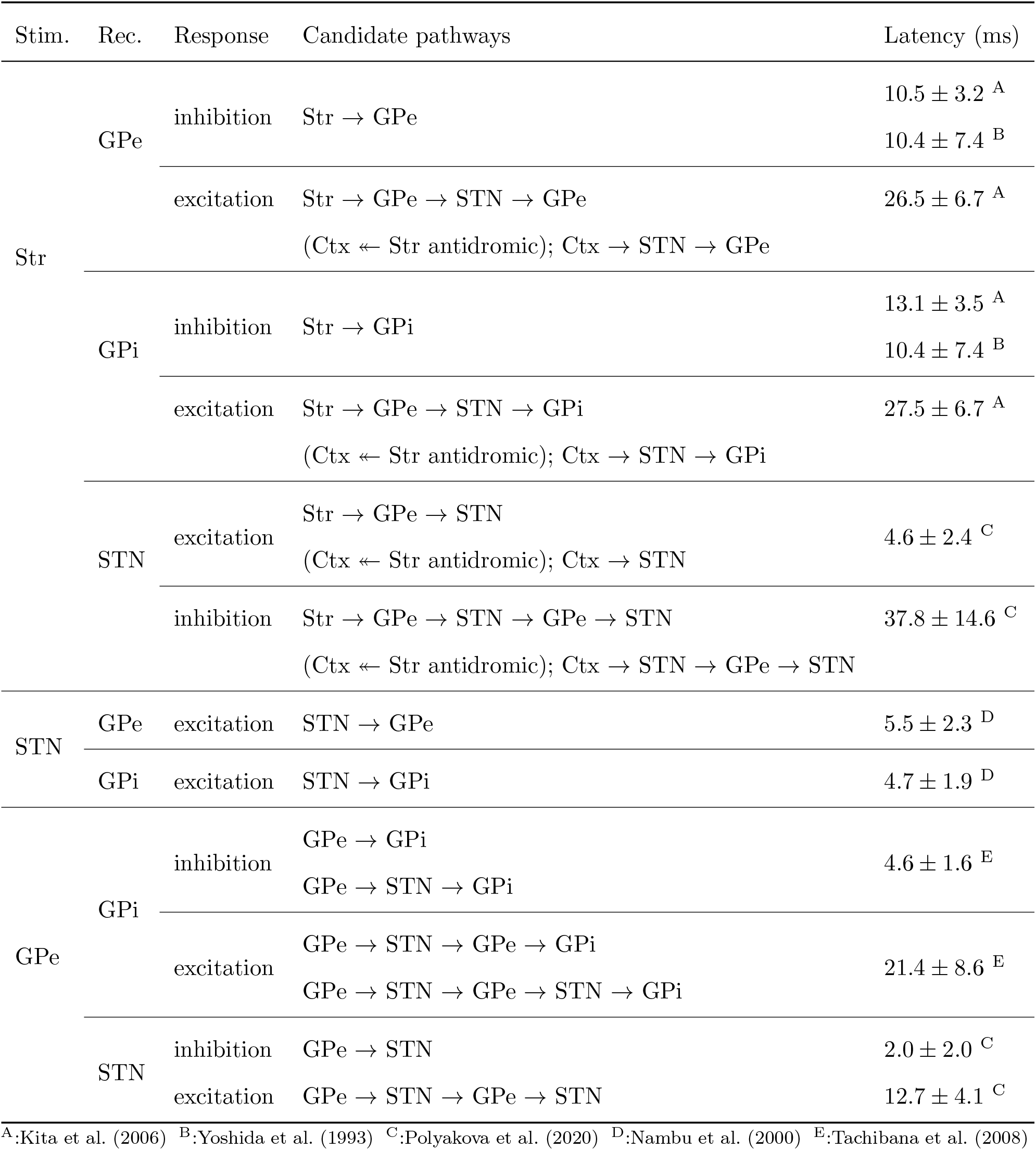
Summary of the candidate pathways and reference data in ms (Part 1). The column “Stim.” indicates the location of the stimulation; “Rec.” indicates the location of the recording; “Response” indicates the type of response recorded; the candidate pathways along the quantitative data expressed as mean*±*SD constitute the remainder of the table. In the case of striatal stimulation, we also consider pathways involving antidromic excitation of cortical axons, noted as “Ctx ↞ Str antidromic” (see methods for details).

The second effect, i.e. the STN excitation following a striatal stimulation in less than 5 ms, seems to be in contradiction with previous data. The shortest pathway supporting this effect within the basal ganglia is the Str → GPe → STN one. But the results of Kita et al. (2006) and Yoshida et al. (1993) suggest that the Str → GPe pathway has an average latency of 10ms, implying that the Str → GPe → STN pathway should be longer than that, and are thus not compatible with a twice smaller total latency. A pathway through the thalamus, not included in the search we performed, would not solve the apparent contradiction: it would be a Str → GPi → Th → STN pathway, that goes trough the Str → GPi connection, whose latency (Kita et al., 2006, Yoshida et al., 1993) is also larger than 10 ms.

This first set of delays thus appears to be the best possible compromise (*Cmpr* column in Table 1) to reconcile these two new short-latency events presented in (Polyakova et al., 2020) with the previously available data, but is yet not fully satisfactory.

One possibility, to solve the apparent contradiction concerning the early Str→ STN excitation, would be to find another pathway to handle this effect. We explored the possibility that this observation would result from an antidromic activation of the cortex, that would then excite the STN. Including this new pathway in the search resulted in the *Antdrm* set of delays (Table 1 and Fig. 2, middle). This second set of delays now predicts an early Ctx→ STN inhibition that should be detectable, as it happens 5 ms before the documented late excitation. Moreover, while using this antidromic pathway allows the Str→ STN excitatory event latency to fit nicely in the expected interval, it is also recruited to explain the Str→ STN inhibitory event, the Str→ GPe excitatory event and the Str→ GPi excitatory event, causing these latencies to become quite low (almost two standard deviations away from the averages documented in the experimental data). The introduction of this hypothesis of an antidromic activation is thus not satisfactory, as it solves some problems but simultaneously introduces new ones.

Finally, we searched what would be the best set of transmission delays if we discarded the data from (Polyakova et al., 2020). The resulting set (*NoPlkv* column in Table 1) also predicts a detectable inhibitory event in the STN after cortical stimulation, as it happens 9 ms before the late excitatory event. The other limitations of this third set of delays are: the conjunction of a Ctx→ STN late excitation latency that is a bit too large, while the following final inhibition is a bit too low; Str→ GPe and → GPi excitation latencies that are also in the lower range. On this restricted set of data, it scores better than the *Cmpr* set, but is also not fully satisfactory.

As such, in the remainder of the article, we will both use the *Cmpr* and the *NoPlkv* sets, and thus duplicate all simulations.

Many previous computational studies assumed the STN → GPe and GPe → STN axonal delays to be similar or equal (Table 1). By contrast, all our candidate explanations yield largely different delays for these projections in the three considered delay sets. Surprisingly, the *Cmpr* set and the *NoPlkv* set have opposite tendencies: the STN → GPe is the slowest projection (9 ms) according to the *Cmpr* set, while it is the GPe → STN (7 ms) according to the *NoPlkv* one.

An intuitive explanation for the imbalance of these two delays inside the same loop may be gained by observing the timing of the early excitation of GPe after cortical stimulation (recorded after roughly 9 ms in Nambu et al., 2000, Kita et al., 2004, cf. Table 3). The quick early excitation of GPe is most likely to be conveyed through the Ctx→ STN → GPe pathway, since the alternative pathway Ctx→ Str→ GPe → STN → GPe involves several other loops and is incompatible with other timings. Furthermore, given that the Ctx→ STN connection conveys excitatory events in about 6 to 7 ms (Nambu et al., 2000, Iwamuro et al., 2009, 2017, Polyakova et al., 2020), the STN → GPe connection delay has to be around 2 to 3 ms. Intuitively, such a quick STN → GPe connection implies a slow GPe → STN connection in order to satisfy the other timing constraints within the basal ganglia (Table 3 and 2). For example, the Ctx→ GPe, the Str→ GPe, the Str→ GPi and GPe → GPi late excitatory events all transit through the STN *↔* GPe loop. This is indeed what the *NoPlkv* set predicts. Mechanistically, this delay imbalance is consistent with the potential myelination of glutamatergic STN axons. Corroborative with the *NoPlkv* set, evidence of the existence of STN axons myelination has been reported in monkey and rat studies (Yelnik and Percheron, 1979, Koshimizu et al., 2013). Incidentally, a detailed computational study of STN neurons has shown that their myelination may mediate the therapeutic effects of deep brain stimulation (Bellinger et al., 2008). On the contrary, and as previously mentioned, the *Cmpr* set is heavily constrained by the minimal GPe → STN latency required by the (Polyakova et al., 2020) data, such that the STN → GPe delay is used to compensate and prolong the duration of the transit of the signals inside the STN *↔* GPe loop.

**Table 3:**
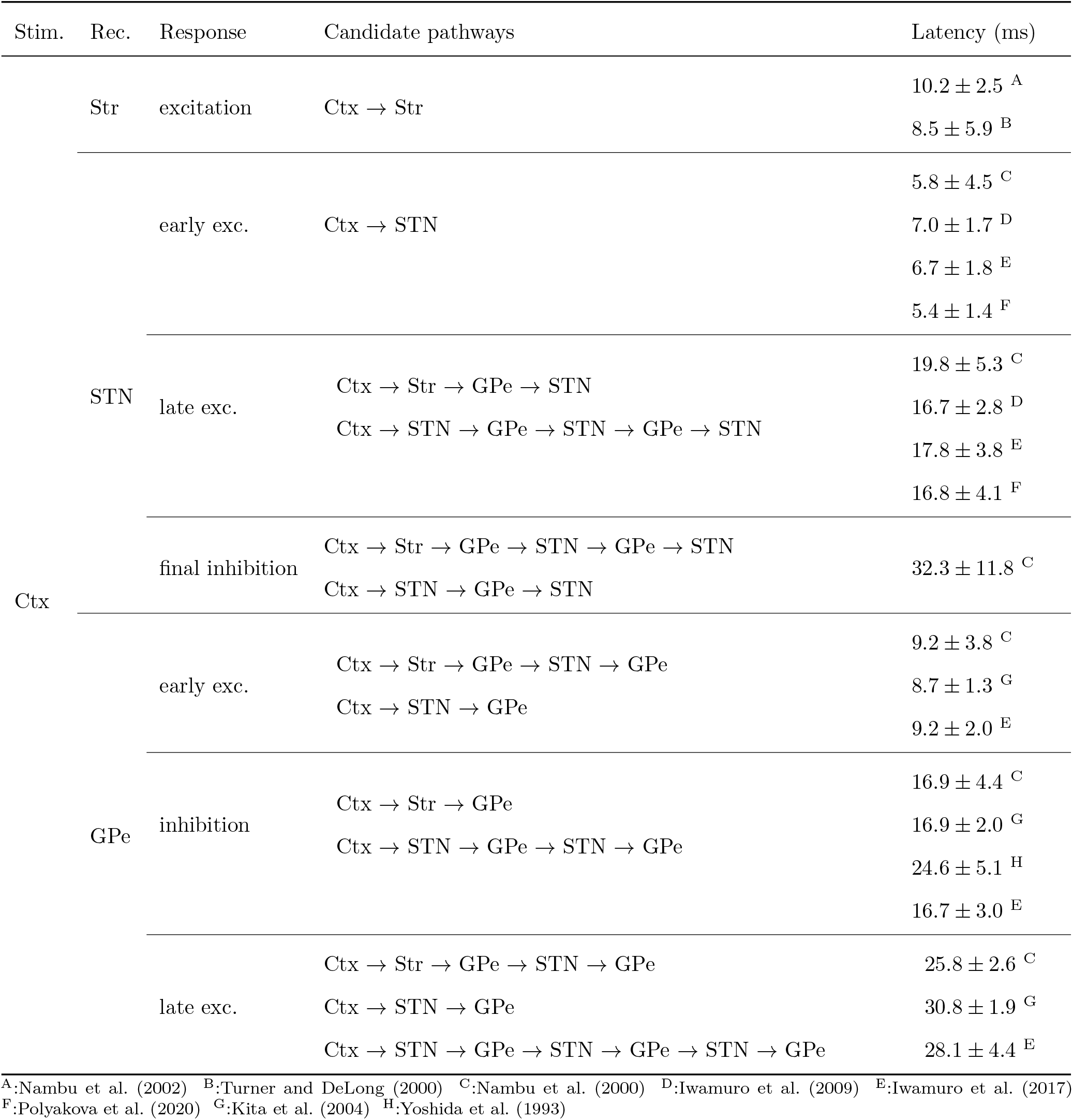
Summary of the candidate pathways and reference data (Part 2). See Table 2 for notations.

### Conditions for the Emergence of *β*-Band Oscillations

We simulated the effect of extra-striatal DA depletion on both the pre-synaptic D2-like receptors in the GPe and the STN, and the post-synaptic D5 receptors in the STN (See Eqs. 10 and 11), while incorporating the axonal delays described in the previous section. Our results show that an increase of the PSP amplitude in the GPe and the STN and a not too strong increase of the mean threshold between resting and firing rates in the GPe and the STN generate an oscillatory regime for both sets of delays (dark-blue regions of the parameter exploration matrices of Figs. 3A and 4A). This oscillatory regime affects all the simulated populations (fig. 5), and because of the fundamental differences in the transmission delays in the STN *↔* GPe loop for the two considered sets of delays, the STN peak activity precedes the GPe one (fig. 5, “GPe & STN” panels) with a longer duration with the *Cmpr* one (11 ms) than with the *NoPlkv* one (3 ms). The frequency of these oscillations are in the upper *β*-band, with an average frequency of 32Hz for the *Cmpr* set of delays, and 35Hz for the *NoPlkv* one (panels B of figs. 3 and 4). It is interesting to note that in the normal regime of operation (power spectra in figs. 3C and 4C), the tendency of the model to favour these frequencies is already visible.

**Figure 3:**
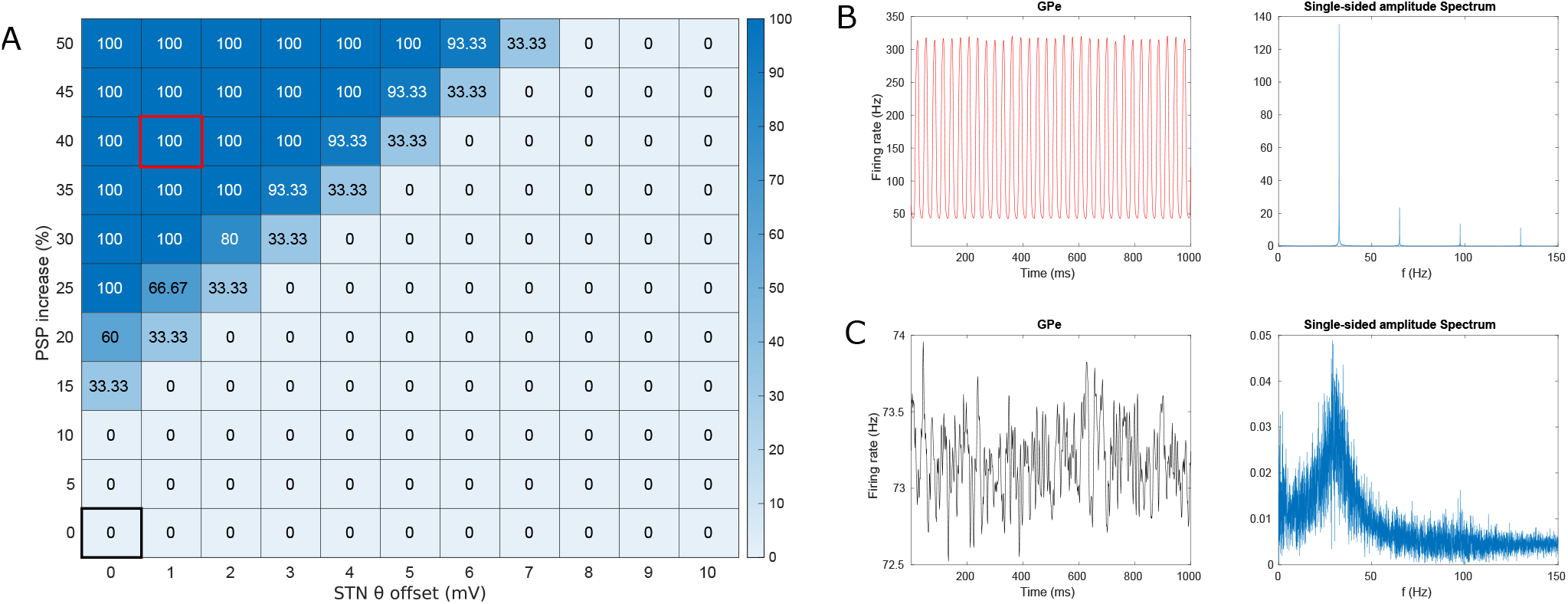
Emergence of oscillations under dopamine depletion with the *Cmpr* set of delays. A: proportion of the basal ganglia models that oscillate depending on the increase in the PSPs in the GPe and STN (D2 receptors) and on the increase in the firing threshold of the STN (D5 receptors). B: Oscillatory GPe activity (left) and the corresponding power spectrum (right) corresponding to a PSP increase of 40% and a STN threshold increase of 1 mV (shown with a red square in A). C: Irregular GPe activity (left) and the corresponding power spectrum (right) corresponding to the model without dopamine depletion (shown with a black square in A).

**Figure 4:**
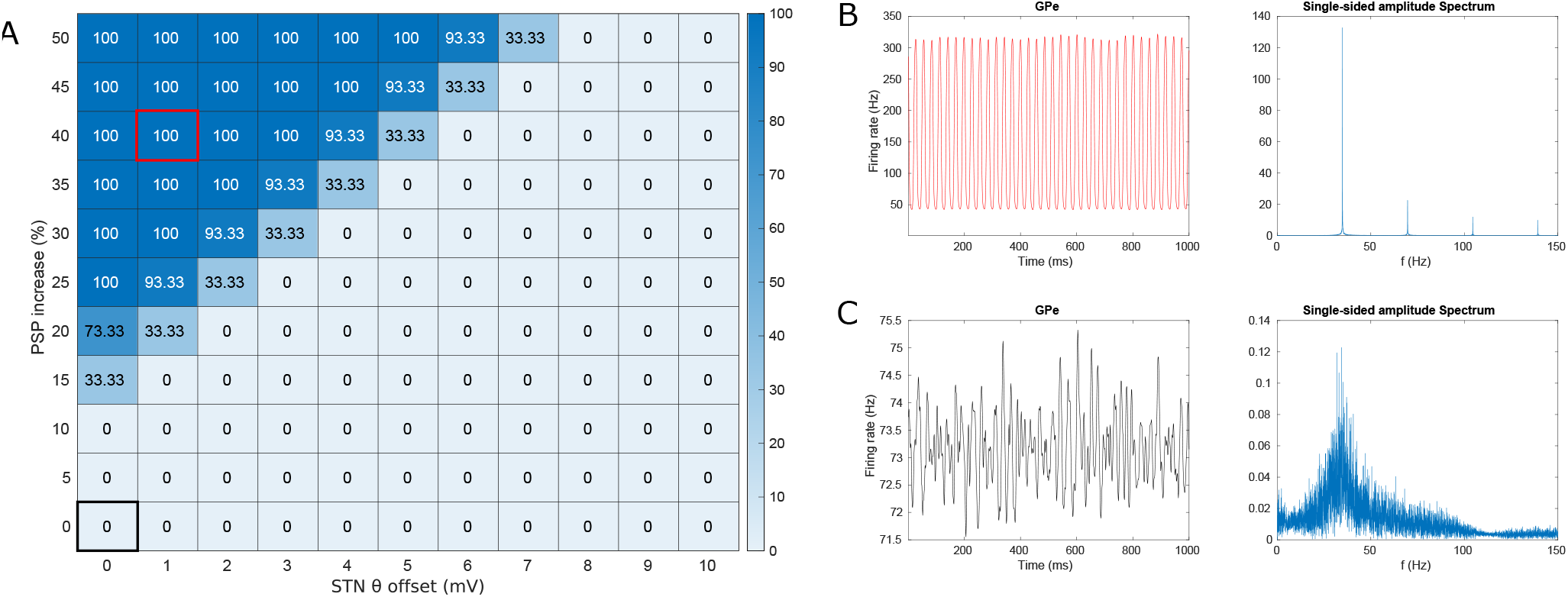
Emergence of oscillations under dopamine depletion with the *NoPlkv* set of delays. A: proportion of the basal ganglia models that oscillate depending on the increase in the PSPs in the GPe and STN (D2 receptors) and on the increase in the firing threshold of the STN (D5 receptors). B: Oscillatory GPe activity (left) and the corresponding power spectrum (right) corresponding to a PSP increase of 40% and a STN threshold increase of 1 mV (shown with a red square in A). C: Irregular GPe activity (left) and the corresponding power spectrum (right) corresponding to the model without dopamine depletion (shown with a black square in A).

**Figure 5:**
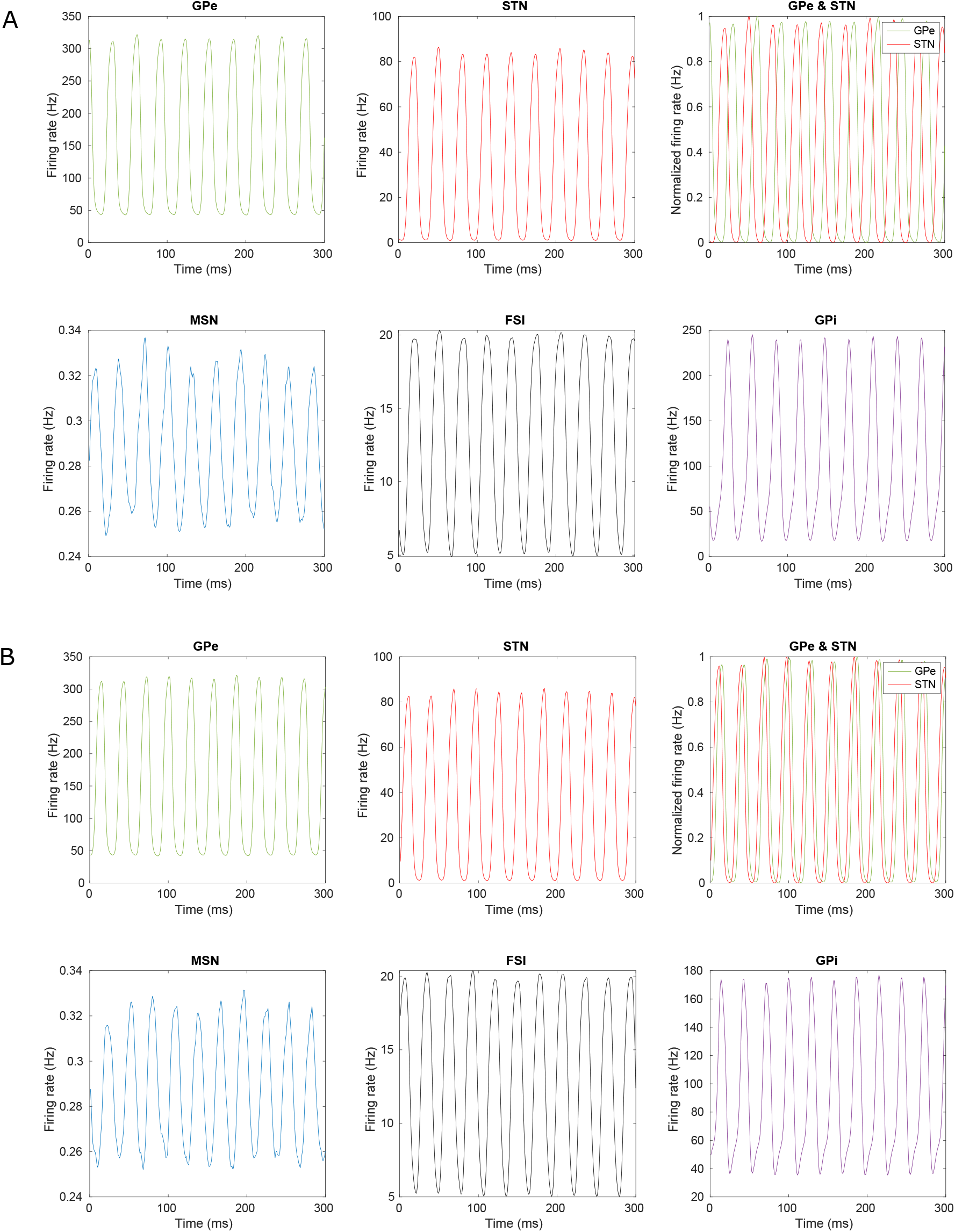
Oscillatory activity in the whole basal ganglia circuit, caused by DA depletion. These illustrative simulations use the parameterization #2 of the BCBG, a GPe and STN PSP increase of 40% and a STN firing threshold increase of 1mV. A: with the *Cmpr* set of delays; B: with the *NoPlkv* set of delays

The conditions of generation of a high-*β* oscillatory behaviour in the STN and the GPe, when simulating extra-striate dopaminergic depletion, require a trade-off between the facilitation of the PSP, driven by the D2 receptors and the weakening of the STN excitability by the D5 receptors.

It is worth noting that although our simulations also reported oscillatory activity following DA depletion in other nuclei such as the GPi and Striatum, including MSN and FSI cells (Fig. 5), these oscillations did not originate locally. Replacing the simulated pallido-striatal and subthalamo-striatal activities with synthetic signals mimicking the baseline obtained in the normal condition, cleared the striatal oscillatory activity (figs. 6 and 7, “no feedback to striatum” panels), thus confirming the GPe *↔* STN origin of the oscillations. By contrast, if the GPe → FSI connection remained active (see Fig. 6 and 7, “GPE → FSI enabled” panels), the oscillations re-emerged in the Striatum. Note also that keeping this feedback connection active, and thus the MSN → GPe → FSI → MSN loop, increased the amplitude of the oscillations in the GPe and the GPi. The GPi provides no input to other basal ganglia nuclei, and thus cannot be involved in the observed emergence of oscillations. In conclusion, oscillatory activity in the Striatum and GPi does not originate locally, but is conveyed from the phase-locked oscillatory activity of the GPe *↔* STN loop.

**Figure 6:**
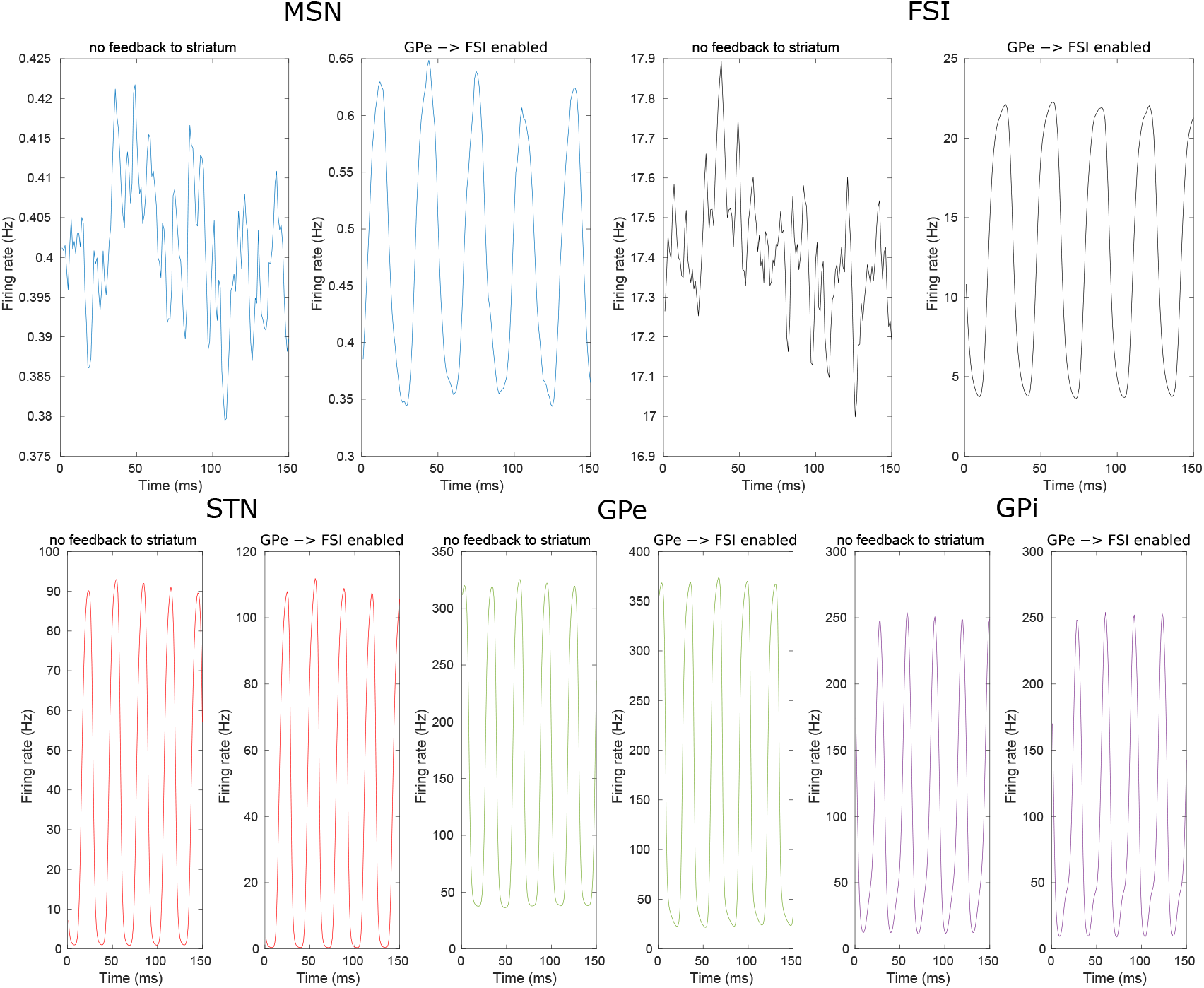
Transmission of the oscillations in the circuit (*Cmpr* set of delays, BCBG model parameterization #2, 40% GPe and STN input PSP increase, 1mV STN threshold increase). In the first set of simulations (left part of each panel), the feedback from the GPe and the STN to the striatum was overridden: STN and GPE input were replaced by synthetic inputs mimicking the normal GPe and STN activities. In the second set of simulations (right part of each panel), the GPe →FSI feedback was selectively re-activated. Oscillations in the MSNs and the FSIs appears only in the second set of simulations, showing that striatal oscillations are byproduct of GPe-STN oscillations mediated selectively through the GPe → FSI connection.

**Figure 7:**
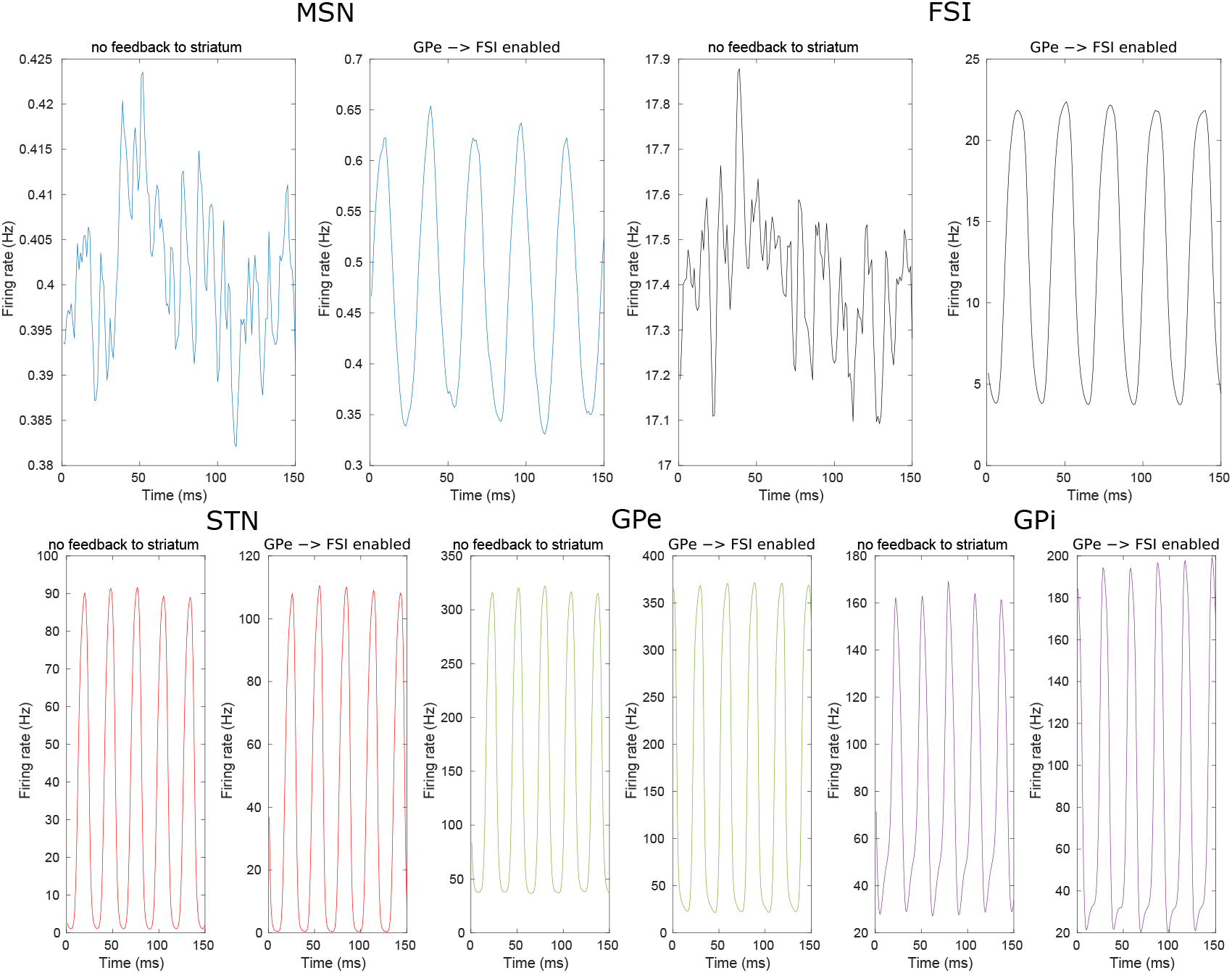
Transmission of the oscillations in the circuit (*NoPlkv* set of delays, BCBG model parameterization #2, 40% GPe and STN input PSP increase, 1mV STN threshold increase). In the first set of simulations (left part of each panel), the feedback from the GPe and the STN to the striatum was overridden: STN and GPE input were replaced by synthetic inputs mimicking the normal GPe and STN activities. In the second set of simulations (right part of each panel), the GPe→FSI feedback was selectively re-activated. Oscillations in the MSNs and the FSIs appears only in the second set of simulations, showing that striatal oscillations are byproduct of GPe-STN oscillations mediated selectively through the GPe → FSI connection.

The frequency of the oscillations depended essentially on the projection delays between the GPe and the STN. We explored, for both sets of delays, the changes on oscillation frequency caused by all possible variations of the GPe → STN and the STN → GPe delays between 1 and 13ms (Fig. 8), while keeping the rest identical: the sum of these delays clearly defines the oscillation frequency.

**Figure 8:**
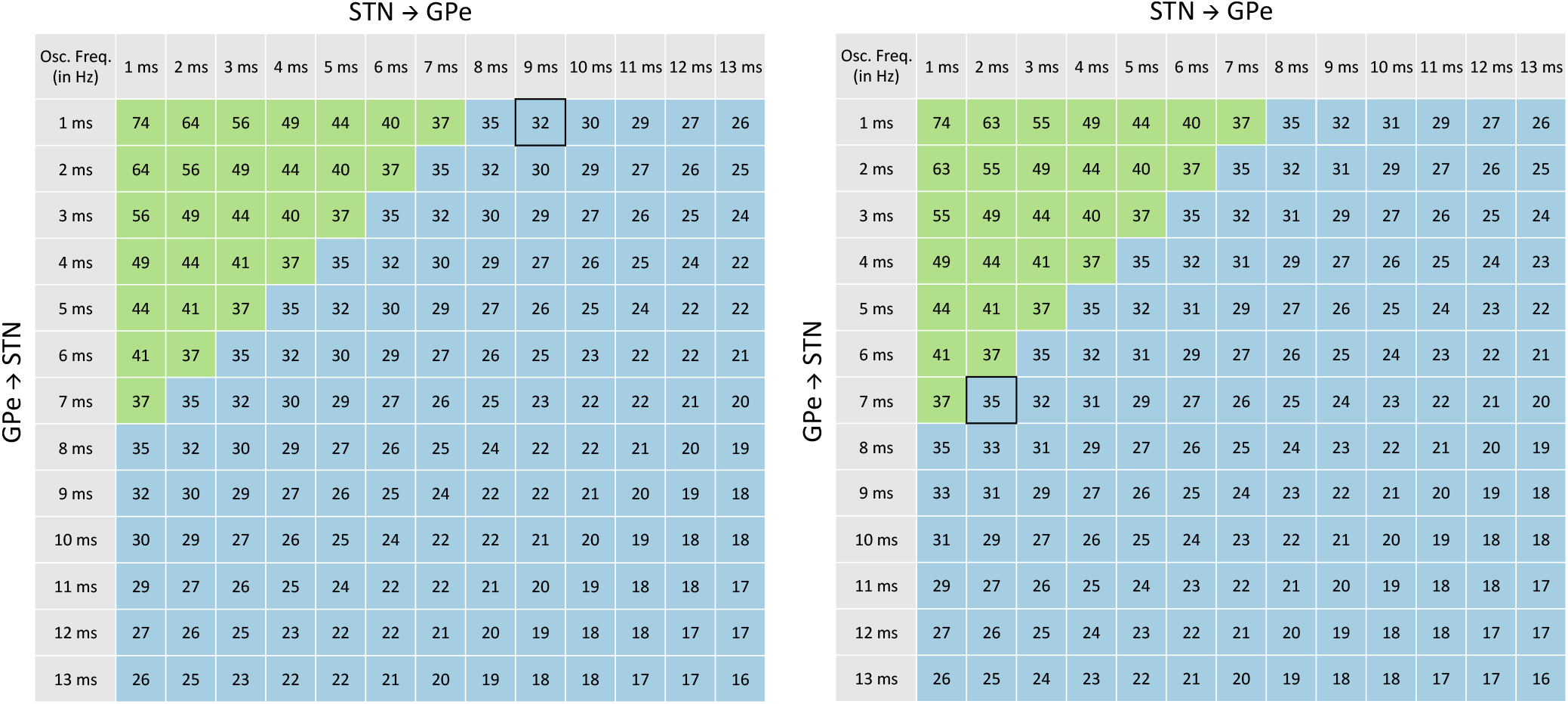
Frequency of oscillation (in Hz) as a function of the axonal delays between STN and GPe. Delays from STN to GPe are listed as columns and from GPe to STN as rows. The other delays are set according to the *Cmpr* set of delays (left) and the *NoPlkv* set of delays (right). The optimized axonal delays are shown in the cells marked with a black rectangle. Colors indicate different oscillation regimes, with the *β*-band (15-35 Hz) shown in blue and the *γ*-band (35-80 Hz) shown in green.

## 3. Discussion

This study has proposed a simple method to combine the existing stimulation experiments used to measure the latency of excitatory and inhibitory events in the basal ganglia, so as to estimate the transmission delays in the various connections of the circuit. This method highlighted some apparent contradictions in the experimental data concerning the GPe to STN projection, which led us to propose two sets of delays, a first one providing the best possible compromise resulting from this data, and a second one ignoring the measures obtained in the Polyakova et al. (2020) study that introduced the potential contradiction (columns *Cmpr* and *NoPlkv* in table 1, respectively).

We have also shown that specific changes of biophysical properties within the GPe-STN loop are sufficient to trigger oscillations in the high *β*-band. These predictions resulted from the introduction of these sets of delays within a computational model of the macaque monkey basal ganglia, fitted from over a hundred independent anatomical and physiological experimental data in macaque monkeys (Liénard and Girard, 2014). Although this model contains numerous parameters, it is important to notice that these were optimized to fit to healthy non-human primate data. By contrast, the results and predictions of this study result from varying two free parameters only, which model the process of DA depletion in GPe and STN: the D2 PSP amplification and D5 firing threshold increase (Figs. 3 and 4). A first prediction of our model is that extra-striate D2 receptors (in the GPe and the STN) and D5 receptors (in the STN) play opposing roles in the generation of *β*-band oscillations when DA decreases: D2 receptors cause them to appear and gradually increase their intensity, and D5 receptors attenuate them. A second prediction is that decreasing the STN D5 receptor activity beyond its constitutive activity (i.e. degrade the D5 receptors normal behaviour even stronger than with DA depletion) would shift the dynamics of the system towards a steady state with no oscillations (see Fig. 3 and 4, shift from the dark-blue to the light-blue area along the STN *θ* offset axis). This is consistent with observations by Chetrit et al. (2013), who showed that diminishing the D5 receptor constitutive activity in the STN of 6-OHDA PD rats did cancel abnormal neuronal activity and reversed motor impairment. Finally, the oscillatory activity reported in other nuclei (e.g. within the FSI-MSN circuitry) does not originate locally, but is rather relayed by projections from the GPe/STN loop (figs. 6 and 7).

### Axonal delays

Numerous stimulation studies participated in characterizing transmission delays across different neuronal groups of the BG circuitry (Yoshida et al., 1993, Turner and DeLong, 2000, Nambu et al., 2000, 2002, Kita et al., 2004, 2005, 2006, Tachibana et al., 2008, Iwamuro et al., 2009, 2017, Polyakova et al., 2020), to the best of our knowledge, this work is the first to combine these experimental studies by computational means in order to determine the set of delays that could be compatible with all of them. In a review dedicated to the GPe, Jaeger and Kita (2011) proposed in their Fig. 1 a summary of the results obtained by the then-available stimulation studies, in which a set of transmission delays was proposed to explain the latencies of excitatory and inhibitory events (panel A of their fig. 1). The details of the method used to propose this set of delays was unfortunately not documented in the main text. We have evaluated the score of this set with our method (Fig. 9), it clearly performs worse than the sets reported in this manuscript. In particular, it tends to underestimate the latency of all the late effects. Interestingly, this set of delays also predicts an early inhibition in the STN after cortical stimulation (with a 10 ms latency), before the late excitation and the final inhibition. In the panel B of their Fig. 1, they suggest that this inhibition would be sufficient to stop the STN early excitation, but may not be strong enough to generate an inhibitory event *per se* (i.e. a decrease of the activity below the baseline, strong enough and long enough to be categorised as an inhibitory event), explaining why its is not reported in experimental studies.

**Figure 9:**
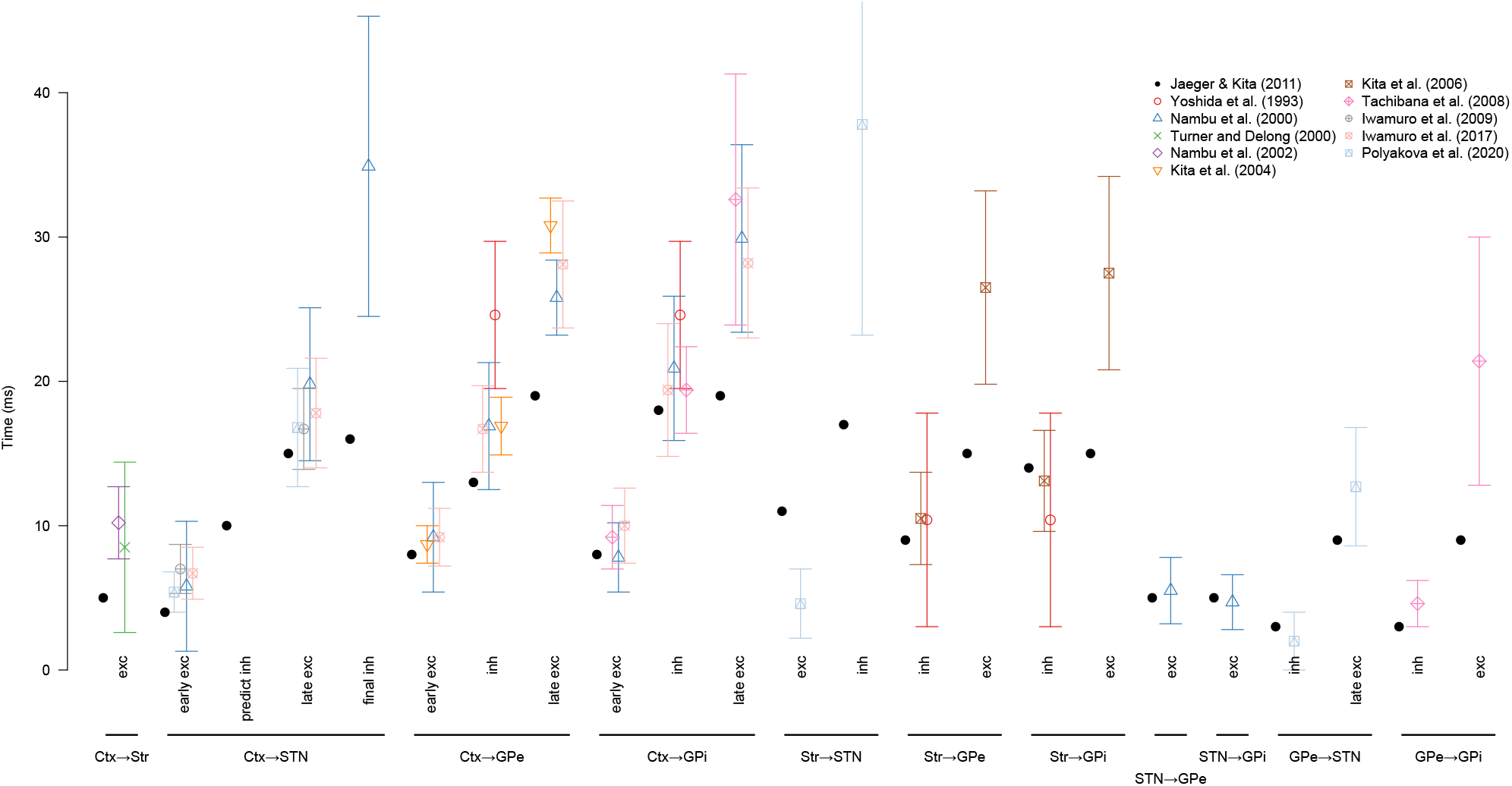
Comparison of the latencies of the excitatory and inhibitory effects measured after the stimulations (averages and standard deviations) from our set of reference studies, with those predicted by the set of delays of (Jaeger and Kita, 2011)

As mentioned above, the results from (Polyakova et al., 2020) raise questions that require further investigation on both the experimental and the modeling sides. First, the extremely short transmission delay from the GPe to the STN (1 ms) measured in this study imposes, in order to stay as much as possible coherent with the rest of the data, a STN to GPe delay (9 ms) that is much larger than what suggests the Nambu et al. (2000) direct measure (5.5 *±* 2.3 ms). Second, the 5 ms delay between a striatal stimulation and an excitatory event in the STN does not appear to be compatible with the most obvious and shortest pathways for its transmission (Str→ GPe → STN or Str→ GPi → Th → STN), as they recruit projections whose transmission delay is at least 10 ms (Str→ GPe or Str→ GPi). However, the apparent inconsistency of these measures may purely result from the limitations of our methodology: we adopted a very crude way of estimating the total duration of the transmission of a stimulation along a pathway (summing transmission delays with a neural processing delay common to all BG neural populations), which does not honour the real complexity of the dynamics of the basal ganglia neural circuit, and we cannot also exclude the possibility that we neglected possible alternate transmission pathways, especially those that would transit outside the basal ganglia proper. A more subtle approach to model these phenomena may thus help solving these paradoxes. Nevertheless, an experimental replication of the STN recordings following striatal and GPe stimulations would be quite useful to acertain that the sources of these potential inconsistency come from the model.

### Striatal and extra-striatal contributions to pathology

Our results suggest that *β*-band oscillations may arise in the BG circuitry when DA is scarce, irrespective of the effect of DA depletion onto DA receptors in the MSN and the FSI in the Striatum, which our model deliberately excludes. Although the extent to which Striatal cells contribute to these oscillations remains to be addressed, our results show that the direct/indirect pathway segregation may be neither central nor necessary to understand the PD oscillatory phenomena in primates.

By removing our focus from the segregation of striatal D1/D2 receptors, we implicitly assumed that the excitatory and inhibitory effects of DA in the striatum cancel each other, yielding no effect on MSN firing rate in average. Consistently with this, electrophysiological studies in primates reported no change in the striatal firing rate after MPTP injection (Goldberg et al., 2002). Furthermore, the modification of the distribution of the firing rates in the MSN population (decreased activity in D1 neurons, increased activity in D2 ones) would exert little influence on the STN *↔* GPe loop, as a consequence of the massively collateralized striato-fugal projection in primates (Parent et al., 1995, Lévesque and Parent, 2005, Nadjar et al., 2006).

Finally, we did not model the effect of DA depletion on the GPi, first, because experimental data on the effects of such depletion is incomplete and contradictory effects have been reported (Rommelfanger and Wichmann, 2010), and second, because the GPi projects only outside the modeled BG circuitry, and thus cannot participate in the local generation of oscillations. Should the model be extended to encompass the cortico-baso-thalamo loops, the GPi would become a central actor of these loops, and as such would participate actively in their dynamics. Modeling GPi DA receptor would then become essential to determine the effects of DA depletion across the circuit.

The expression of D5 receptors in the GPe, similar to what is found in the STN, and their effect on neural activity is unclear (Rommelfanger and Wichmann, 2010). Thus, for the sake of completeness, we simulated the same increase of the firing threshold in the GPe as in the STN. Adding that modulation only marginally modifies the reported boundaries of the oscillation region in the parameter space (fig. 10, A and C). Indeed, isolating the effect of these putative GPe receptors by removing the D5 modulation in the STN revealed that increasing the firing threshold of the GPe does not really affect the emergence of oscillations. This suggests that cancelling the constitutive activity of D5 receptors in the GPe should not replicate the electrophysiological and motor observations Chetrit et al. (2013) obtained in the STN.

**Figure 10:**
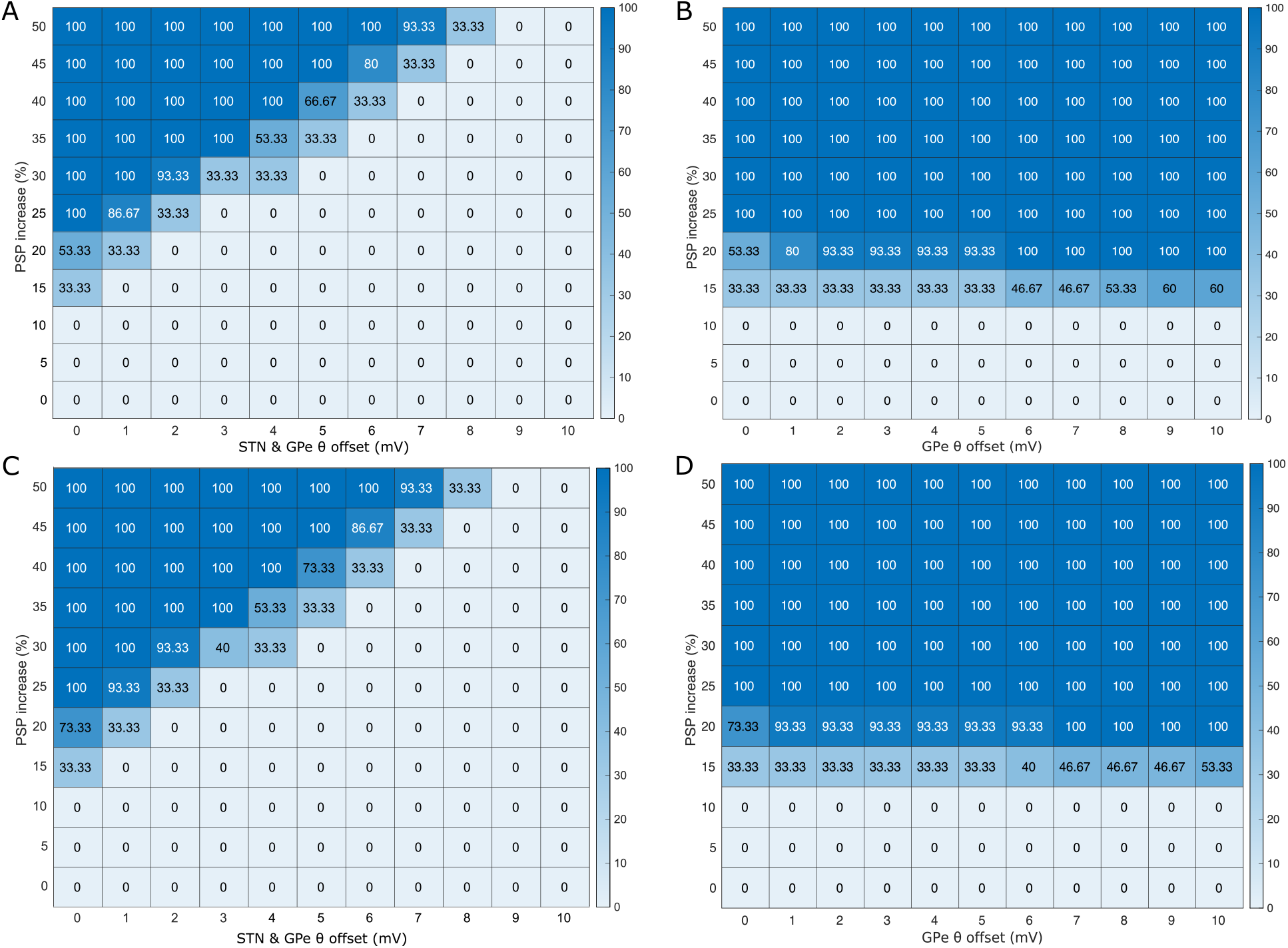
Effect of the simulation of D5 receptors in the GPe with regards to the emergence of oscillations under dopamine depletion. A, B: using the *Cmpr* set of delays; C, D: using the *NoPlkv* set of delays. Same as figure 3A, except that in A and C, *θ* is increased in both STN and GPe, and, in B and D, only in the GPe.

A clear limitation of the model is that the oscillations it generates are in the highest part of the *β*-band, while MPTP monkeys rather exhibit oscillations in the lower range (Tachibana et al., 2011). This may result from a number of simplifications of our model. First, the dynamics of the simulated neural populations depend exclusively on the dynamics of the post-synaptic potentials at the synaptic level, no other internal dynamics of the neuron itself is simulated, that could dampen the oscillations. Second, no synaptic adaptation mechanisms were included, while (Shouno et al., 2017) suggest that short-term facilitation and depression participate in the GPe-STN loop behaviour. Investigating whether these simplifications are sufficient to explain this discrepancy is the matter of future work.

### Theories of *β*-band Oscillations

There are several theories for the origin of *β*-band oscillations in the BG: the *striatal origin theory*, based on the hypothesis of changes of intrinsic properties of striatal MSNs (McCarthy et al., 2011); the *striatal inhibition theory* on the increased striatal inhibition by means of the D2 neurons on the GPe-STN loop (Kumar et al., 2011, Lindahl and Hellgren Kotaleski, 2016); and the *FSI loop theory*, which posits that (in mice) the GPe to Striatum feedback projection (through the FSI) is the cause of the oscillations (Corbit et al., 2016). When it comes to explain primate data, these theories have been nevertheless challenged by experimental evidence showing that the GPe still exhibits oscillatory behaviour after having severed its inhibitory inputs from MSN (Tachibana et al., 2011), and by the absence of segregated striato-fugal pathways. Concerning the *FSI loop theory*, our results show that, in primates, the role of the MSN-GPe-FSI loop is consistent with a relay transferring GPe-STN oscillations back to the striatum (see Figs. 6 and 7), and that it is unlikely to be a source. Additionally, Van Albada and Robinson (2009), Van Albada et al. (2009) introduced the hypothesis of the *thalamo-cortical loop* being the primary cause of oscillation. Again, this hypothesis seems unlikely, as (1) it requires the segregation of direct/indirect pathways which is non-existent or partial in primates, and (2) it is severely constrained by long intrinsic delays, which are more suitable to sustain *θ* than *β*-band oscillations (Leblois et al., 2006b). Finally, Pavlides et al. (2015) showed that it is possible for a properly parametrized cortico-basal model encompassing three loops (the STN *↔* GPe, the intra-cortical and the cortico-basal) to contribute to the emergence of *β* oscillations, without any of them being capable of autonomously sustaining them.

By contrast to the aforementioned hypotheses and models show that the STN *↔* GPe loop of primates is ready to oscillate autonomously in the *β*-band by simply increasing the coupling between both nuclei. Furthermore, although previous research had already hinted this loop to be the source of sustained *β*-band oscillations (Gillies et al., 2002, Terman et al., 2002), we are first to pinpoint the opposing roles of D2 and D5 extrastriate receptors in the emergence of these oscillations.

Together with our results, three other independent studies had suggested the GPe-STN loop to be crucial to generate oscillations in the *β*-band. However, they exhibit some degree of inconsistency with experimental data, either in their assumptions or in their implementation, or rely on additional mechanisms not required in our parsimonious model. First, Tsirogiannis et al. (2010) used unrealistic brief transmission delays (1 ms for both GPe → STN and STN → GPe connections, cf. Table 1), assumed PD to influence the time-course of post-synaptic potentials, and relied on the existence of segregated direct/indirect pathways. Likewise, Nevado Holgado et al. (2010) stressed the importance of the transmission delays in this loop to set the oscillation frequency. However, they assumed PD to yield an unrealistically strong change of synaptic strength (two-fold increase for GPe → GPe, almost four-fold for cortex → STN, almost ten-fold for striatum → GPe and GPe → STN). Finally, in their recent spiking model, Shouno et al. (2017) showed that STN-GPe oscillations could emerge if post-inhibitory rebound excitation at the STN level and short-term plasticity in both STN and GPe were introduced. Our model shows that these mechanisms are nevertheless not required.

Humphries et al. (2006) have shown that GPe-STN oscillations in rats would yield *γ*-band and slow (*<* 1Hz) oscillations. To assess the generality of our hypothesis, we transferred the delays of Humphries et al. (2006) to our primate model, as to test whether the predicted frequency of our model would also match data obtained experimentally on that species. Interestingly, when changing the transmission delays of our model to those of the rodent literature (namely 4ms for GPe → STN and 2ms for STN → GPe), our predicted oscillation frequency shifted to 44 Hz (Fig. 8), a value reasonably close to those recorded in rats (53-55 Hz). Although cautiously, this may suggest a common principle of oscillation across both species. Note however that experimental results tend to suggest that the mechanisms of abnormal oscillations in PD models can be different from one species to another: the involvement of the STN reported in monkeys (Tachibana et al., 2011) hasn’t been found in mice (de la Crompe et al., 2020).

## 4. Methods

### 4.1 Characterization of Delays between Primate Basal Ganglia Nuclei

For our model to make sensible predictions about the naturally occurring frequencies of oscillation across basal ganglia nuclei, it was first required to establish the typical transmission delays across them for the case of the macaque brain, which are not directly available. Instead, we had to derive them from the latencies between excitatory and inhibitory events recorded during stimulation studies (Yoshida et al., 1993, Turner and DeLong, 2000, Nambu et al., 2000, 2002, Kita et al., 2004, 2006, Tachibana et al., 2008, Iwamuro et al., 2009, 2017, Polyakova et al., 2020). Therefore, it was first necessary to develop a methodology to properly identify the combinations of pathways and transmission delays involved in each experimental data set (see Tables 2, 3 and 4), so as to extract the specific delays across basal ganglia nuclei.

**Table 4:**
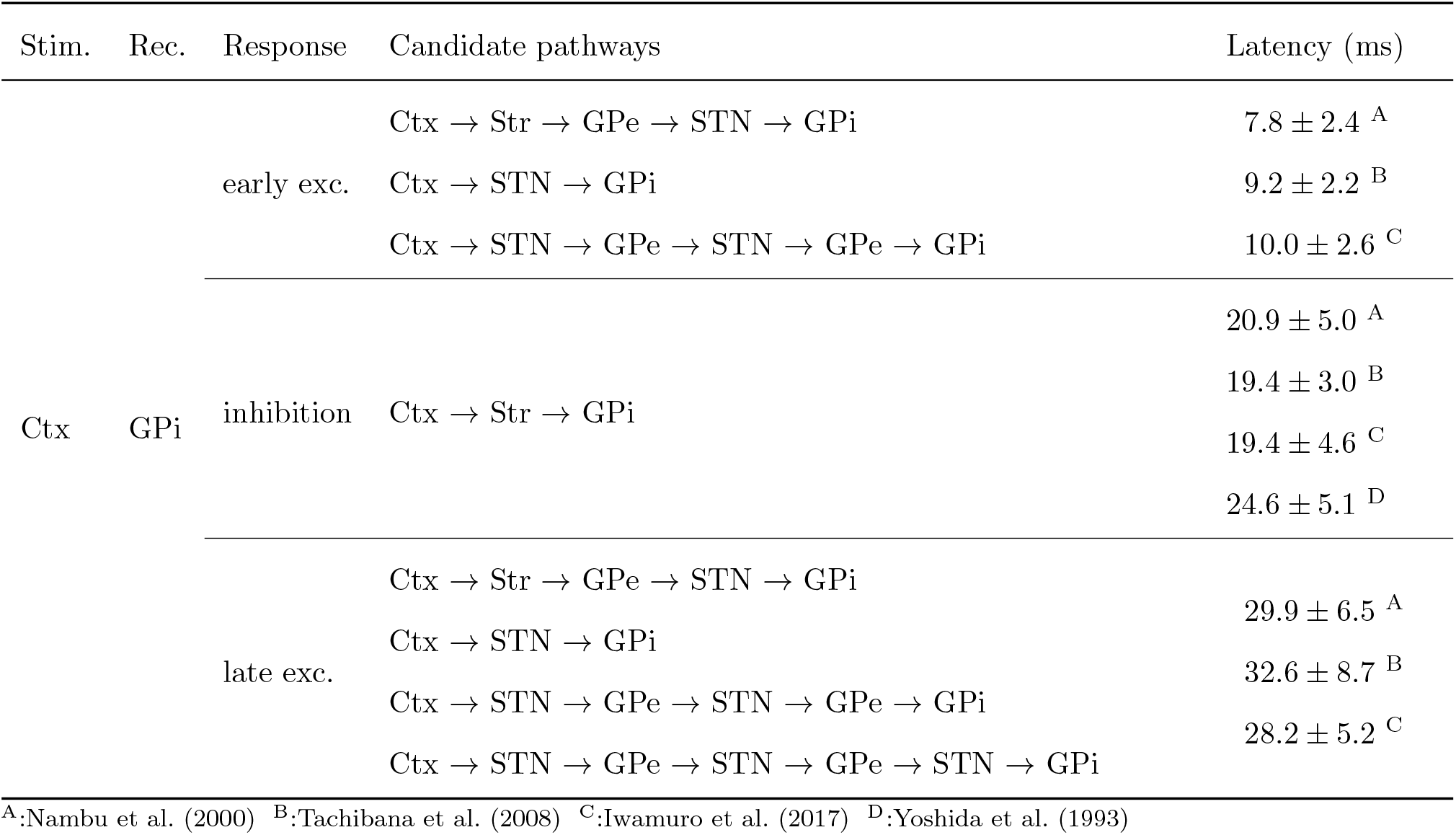
Summary of the candidate pathways and reference data (Part 3). See Table 2 for notations.

To that end, we considered all known connections within the BG (Fig. 11A), with the exception of the sparse projections from STN to Striatum and from GPe to Striatum (Jaeger and Kita, 2011). Indeed, STN stimulation fails to elicit MSN activity (Kita et al., 2005); and although cortical stimulation elicits MSN overactivity, this is not followed by a noticeable second excitation which would have signaled an rebound mediated by the STN (Nambu et al., 2002). Also, if the GPe to Striatum projection were functionally active in stimulation studies, we would expect the subsequent inhibition of the GPe to cause MSN overactivity, an event which has not been observed experimentally (Nambu et al., 2002). Notice that the resulting simplification of the BG connectivity graph is also in line with the results of our previous study (Liénard and Girard, 2014) which showed that the influence of these pathways on the striatum could only be potent if they targeted the FSI. Hence, they could only influence the MSN by silencing them through interneurons, and as MSN are already mostly silent at rest, this should not have observable consequences on the other nuclei.

**Figure 11:**
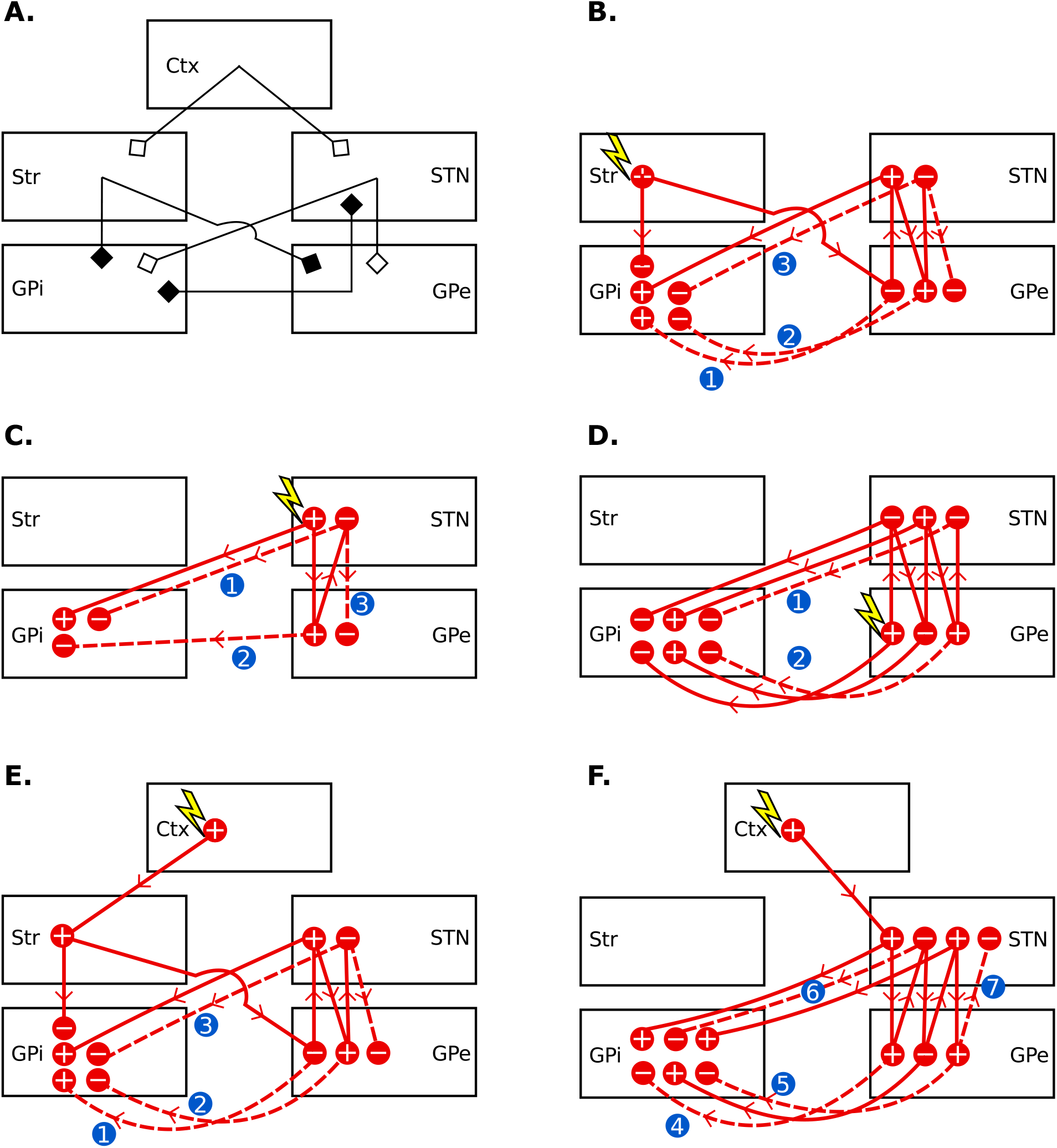
A. Connections considered in the transmission delay estimation. Black endings denote inhibitory connections and white endings denote excitatory connections. B-F. Possible timecourses of a stimulation originating in the striatum (B), STN (C), GPe (D) and cortex (E-F). For clarity, the cortical stimulation is subdivided into one representation of the striatal excitation consequences (E) and of the subthalamic excitation consequences (F). In all panels, circled “+” indicate that overactive nuclei and “-” indicate underactive nuclei. Successive states of overactivity or underactivity are placed from left to right if they can be ordered (e.g. after an excitation in the Str, the STN will always be overactive before being underactive). Successive states that can not be ordered are placed on different lines (e.g. after an excitation in the Str, the first state of GPi could *a priori* be either an underactivity or an overactivity). Dashed numbered links correspond to pathways that are not recruited, see text for their individual justifications.

To match the timings between excitatory and inhibitory events reported in the literature, we first deployed the graph of the projections potentially involved into series of candidate pathways (Fig. 11 discussed in next section). We further trimmed down the number of pathways by excluding highly recurrent ones as to lower the computational complexity of the search. We limit in particular the number of iterations through the STN*↔*GPe loop to two, as more iterations would result in latencies that are too long; other exclusions are discussed in details in section 4.1.1. The resulting set of candidate pathways that are summarized along with the experimental timing references in Tables 2, 3 and 4. Note that some of the studies we include here stimulated different cortical areas, for example Nambu et al. (2000) stimulated the primary motor area (M1) as well as the supplementary motor area (SMA). In such cases, we rely on the shortest latency reported per study.

We then perform an exhaustive exploration of the space of axonal delays to find the best fit. In order to achieve this, we compute the time that would be needed by each pathway in a simplified model to match to the experimentally recorded delay, and score the fit of the quickest pathway with experimental data. To aid the optimization process, all delays were constrained to assume biologically plausible values (1-12 ms). Our exhaustive search to find the optimal fit thus implied the evaluation of all the 12^8^ (*≈* 430 million) possible combinations. Each combination was assigned a score dependent on the amount of experimental data it was able to replicate.

#### 4.1.1 Pathways involved in the stimulation experiments

Stimulations in the cortex, Str, STN or GPe result in an intricate superposition of excitatory and inhibitory effects that could be supported by a multitude of different pathways. We could however simplify the connection graph to rule out several pathways. We will successively review the different stimulation locations, the excitatory or inhibitory responses that they cause and the pathways that could be mediating these responses (Fig 11A).

First we considered the case of the striatal stimulation (Fig. 11B). Following this stimulation, GPe and GPi neurons are first inhibited, then excited (Kita et al., 2006). The artificial blockade of STN eliminates the excitation but does not affect the inhibition (Kita et al., 2006), hence the STN is required for the excitation. The excitation can thus not possibly be mediated by the (Str→ GPe → GPi) chain because it does not involve the required STN, so we can rule out pathway number 1 of Fig. 11B. Furthermore, the artificial blockade of STN does not change the timing of the inhibition, so we can also rule out the pathways number 2 and 3 of Fig. 11B because the chains (Str→ GPe → STN → GPe → GPi) and (Str→ GPe → STN → GPe → STN → GPe → GPi) involve the STN and result in an inhibition of GPi.

Next we considered a stimulation in the STN (Fig. 11C). Following this, more than 80% of the responding neurons in the globus pallidus are excited (Nambu et al., 2000). As this excitation is not followed by an observable inhibition, we can rule out the pathways numbered 1, 2 and 3 of Fig. 11C because they would lead to an underactivity either in GPe or GPi.

We then considered the GPe stimulation (Fig. 11D). In this experiment, only half of the GPi neurons show an early excitation occurring fast (3.4 *±* 0.9 ms), and they all show later an inhibition that is followed by an excitation (Tachibana et al., 2008). This early excitation can hardly be explained by the basal ganglia pathways, because the chains finishing earliest in the GPi, i.e. (GPe → STN → GPi) and (GPe → GPi), result in an inhibition. The (GPe → Str→ GPi) chain could possibly account for this early excitation, however we did not include the GPe → MSN pathway as it does not seem to be involved in these stimulation experiments (c.f. the beginning of this section), and furthermore the latency of this excitation is clearly too fast to be mediated through the slow Str→ GPi connection. Finally, as discussed in Tachibana et al. (2008), the early excitation could be mediated by STN axons targeting both the GPe and GPi. As this does account for the fact that only half of GPi neurons respond and as other pathways can not plausibly explain an early excitation that is this fast, we will not consider further this early excitation. We can also rule out the pathways numbered 1 and 2 because they suppose a late inhibition following the reported inhibition and excitation in GPi, and Tachibana et al. (2008) did not report such a late inhibition.

Finally, Fig. 11E-F illustrates the case of the *cortical stimulation*. This stimulation leads to three distinct temporal responses in STN, GPe and GPi (Nambu et al., 2000): an early excitation followed by an inhibition, and finally a late excitation. To understand the possible timecourses of this cortical stimulation, we subdivided it according to the chains beginning with the direct striatal excitation (Fig. 11E) and with the direct subthalamic excitation (Fig. 11F). After the artificial blockade of activity in the STN, Nambu et al. (2000) reported that the GPi does not exhibit the early or late excitation. We can thus deduce that the STN is required for these excitations, so we can rule out the pathway 1 of Fig. 11E corresponding to the chain (Ctx→ Str→ GPe → GPi) because the STN is not involved in it. Nambu et al. (2000) also reports that after STN blockade, the GPi exhibits the same inhibition, so we deduce that the STN is not part of the chains leading to an inhibition in GPi and rule out the pathways 2 to 6, corresponding to the chains involving the STN and resulting in an underactivity in GPi. No late inhibition has been reported in Nambu et al. (2000), so the pattern of activity “-” then “+” then “-” in the STN is not plausible. We hence rule out the pathway 7.

#### 4.1.2 Optimization of inter-nuclei delays

To compute the time *t* needed for the stimulation in “nucleus 1” to flow over a given chain (nucleus 1 → nucleus 2 → … → nucleus *n*) and to be eventually recorded in “nucleus *n*”, we use a simple formula:

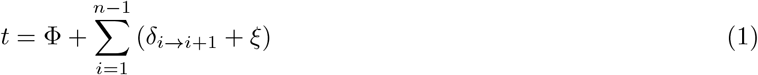

with *δ*_*i→i*+1_ the axonal delay between nuclei *i* and *i* + 1, Φ the time needed for the stimulation to be effective in nucleus 1 and *ξ* the time needed to any nucleus to change their firing rate when receiving the stimulation. The time required for the stimulation to be effectively eliciting action potentials is very small, so we set Φ = 1 ms (a value of Φ = 0 ms was also considered and led to similar results). The time required for the action potential once at the synapse level to be captured by the postsynaptic neuron and to change its potential was considered to be *ξ* = 1 ms, equal for all populations for the sake of simplicity. This latter constant is justified from the shape of the change of intensity after a spike mediated by either AMPA or GABA_A_, because it is already significant after 1 ms (Destexhe et al., 1998) and is in line with the alpha functions that we used in the BCBG model (Liénard and Girard, 2014).

While most pathways modeled involve successive action potentials, we also test separately the existence of cortical antidromic activation of pyramidal tract neurons at striatal stimulation sites. Indeed, such activation of PTN neurons has been observed or hypothesized, resulting then in the direct excitation of the STN through the cortical PTN neurons when stimulating the striatum (Bauswein et al., 1989, Turner and DeLong, 2000). When we model this possibility, we consider that the elicitation of action potentials in cortical neurons is near-instantaneous (Turner and DeLong, 2000) and thus we do not add extra time for its generation. For example, the antidromic pathway that explains the late excitation of GPe after striatal stimulation, noted *(GPe* ↞ *Str antidromic); Ctx→ STN → GPe* in Table 2, has a delay computed as if the stimulation was directly originated from the cortical PTN neurons. Its formula then follows the general shape described by Eq. 1, i.e. *t* = Φ + (*δ*_*Ctx→STN*_ + *ξ*) + (*δ*_*STN→GP e*_ + *ξ*).

As a general rule, each candidate pathway is computed for each response type, and the quickest pathway is assumed to be the one that we observe. The exceptions are the cortical stimulations as they cause an early and a late excitation, in these cases the quickest excitatory response is assumed to correspond to the early excitation and the quickest excitatory response *after the inhibition* is assumed to correspond to the late excitation. Overall, 39 candidate pathways corresponding to 21 couples of (stimulation / recorded response) are checked against 45 experimental data. These experimental data are noted as *T*_*i,j*_ *± σ*_*i,j*_ with *i* the number of the (stimulation/response) pair and *j* the indice of the reference for each pair, as given in Tables 2 and 3. The global score *χ* of the fit between the timing of selected pathways and the reference data is then computed as:

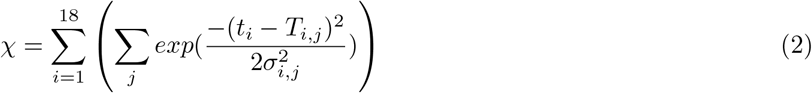

### 4.2 Computational Model of the Basal Ganglia

In this study we focus on the oscillatory activity of the BG circuitry at rest. With the exception of the changes described below (transmission delays optimized to fit to experimental studies and simulation of DA depletion on extra-striate receptors), the model that we present here adopts the mathematical formalism and parameters we previously developed (Liénard and Girard, 2014). Briefly, each nucleus of the basal ganglia is simulated with a mean-field model incorporating the temporal dynamics of neurotransmitters. Inputs from cortical (cortico-striatal neurons, CSN, and pyramidal tract neurons, PTN) as well as thalamic afferents (from the centromedian and the parafascicular nuclei, CM/Pf) are modeled as independent random processes with different average firing rates (Bauswein et al., 1989, Turner and DeLong, 2000, Pasquereau and Turner, 2011, Pasquereau et al., 2015). As in Liénard and Girard (2014), the CSN input was simulated as a Gaussian process centered around 2 Hz, PTN around 15 Hz and CM/Pf around 4Hz. The simulations presented here were obtained using a standard deviation of 2 Hz, corresponding to a high noise in the neuronal activities. The oscillatory patterns obtained with this noise level were similar to those obtained with a lower standard deviation of 0.5 Hz.

The parameter search in Liénard and Girard (2014) extended over more than one thousand optimal model parametrizations that were equally maximizing the plausibility scores defined in that study. An additional assessment showed that the most of variability in these solutions is small jitter (*<* 10^*−*6^ in a search space normalized within [0, 1]) around 15 different base solutions. Thus we restricted our study to these 15 base solutions, as they globally represent the optimal parametrizations of the basal ganglia obtained in Liénard and Girard (2014), as we did in (Girard et al., 2021).

The structure of the model is very close to the one presented in Liénard and Girard (2014), with the exception of the addition of plausible axonal delays and the simulation of dopamine depletion on extrastriate receptors. We use a population model with mean-field formulation. Although we provide here the basic equations of our model, more details about mean-field models can be found elsewhere (e.g. Deco et al., 2008).

One assumption of mean-field models, commonly referred as the *diffusion approximation*, is that every neuron receive the same inputs from another population. We can hence express the mean number of incoming spikes with neurotransmitter *n* per neuron of the population *x* from population *y*:

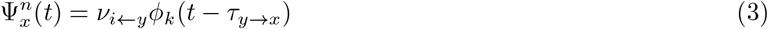

with *ν*_*x←y*_ the mean number of synapse in one neuron of population *x* from axons of population *y, τ*_*y→x*_ the axonal delay between population *y* and *x*, and *ϕ*_*y*_(*t − τ*_*y→x*_) the firing rate of population *y* at time *t − τ*_*y→x*_.

The axonal varicosity counts *ν*_*x←y*_ is the mean count of synapses in population *x* that are targeted by axons from population *y*:

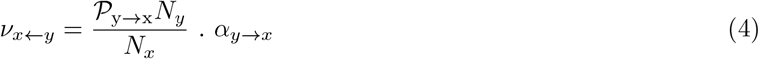

with *N*_*x*_ and *N*_*y*_ the neuron counts of populations *x* and *y, α*_*y→x*_ the mean axonal varicosity count of neurons of *y* with an axon targeting neurons of *x*, and *P*_y→x_ the proportion of such neurons in population *y*.

Mean-field models assume that neurons’ firing thresholds follow a Gaussian distribution. The mean firing rate of a population *x* at time *t* can then be approximated by:

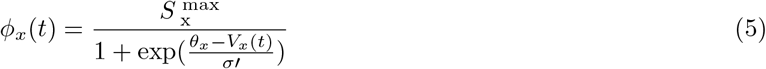

with *V*_*x*_(*t*) the mean potential at the soma at time 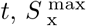 the mean difference between resting and firing thresholds, and, as per Van Albada and Robinson (2009), 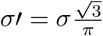 being the standard deviation of the firing thresholds).

The post-synaptic potential (PSP) change to the membrane potential at the location of the synapse contributed by a single spike is modeled by the alpha function (Rall, 1967):

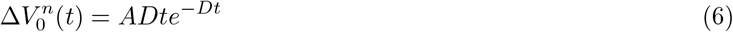

where *A* and *D* relate to the amplitude and duration of PSP and depend on the neurotransmitter *n* mediating the spike. They are computed as follows: *A* = *A*_*n*_ *exp*(1) and 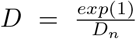 (Tsirogiannis et al., 2010), using the constants reported in Liénard and Girard (2014).

We also model in a simple way the attenuation of distal dendrites as a function of the soma distance. By modeling the dendritic field as a single compartment finite cable with sealed-end boundaries condition (Koch, 2005), we can express for population *x*:

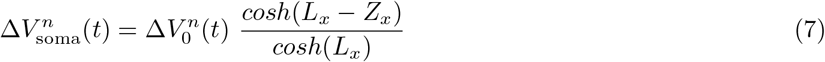

with 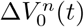 the potential change at the synapse, *L*_*x*_ the electrotonic constant of the neurons and *Z*_*x*_ the mean distance of the synaptic receptors along the dendrites. We further express this mean distance as a percentage of *L*_*x*_: *Z*_*x*_ = *p*_*x*_*L*_*x*_. The electrotonic constant is then calculated according to (Koch, 2005):

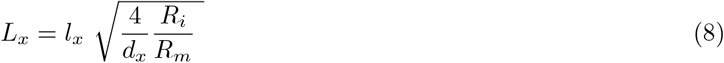

with *R*_*i*_ the intracellular resistivity, *R*_*m*_ the membrane resistance, *l*_*x*_ the mean maximal dendritic length and *d*_*x*_ the mean diameters of the dendrites along their whole extent for population *x*.

The mean potential of the neural population, *V*_*x*_, is finally obtained by integrating the changes of potential caused by incoming spikes over time:

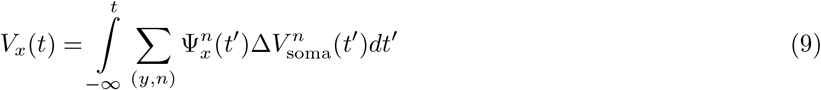

where each couple (*y, n*) represents one afferent population *y* with spikes mediated by the neurotransmitter *n*.

The BG dynamics were simulated with a time-step of 10^*−*4^ ms, as in Liénard and Girard (2014), using a 4^*th*^ order Runge-Kutta integration method.

The code of the model is available on github at https://github.com/SN1885A/BCBG-model.

#### 4.2.1 Extra-striate DA depletion

We hypothesized that an abnormal activation of extra-striatal DA receptors, combined with lagged activity due to inter-nuclei transmission delays, is the primary cause of *β*-band oscillations. To test this hypothesis, we focused on modeling the distribution of DA receptors within the GPe and STN only, as these are the only two nuclei which participate of multiple loops within the BG and could possibly cause oscillations. The GPi was disregarded, as it does not form any closed loop within the BG and thus cannot cause oscillations within the BG.

Since D2 receptors in GPe and STN are located at the pre-synaptic level only (Rommelfanger and Wichmann, 2010), we simulated their deactivation by a dopamine depletion with an increase of post-synaptic potentials following an incoming spike, as follows:

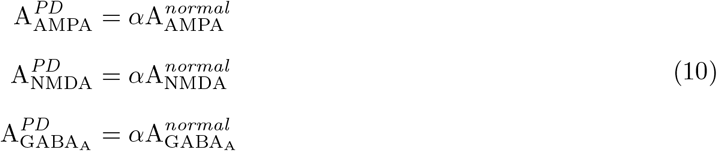

where A_AMPA_, A_NMDA_ and 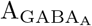 are respectively the peak post-synaptic amplitude of a spike mediated by AMPA, NMDA and GABA_A_. ^*normal*^ denotes their reference value defined in Liénard and Girard (2014), and ^*PD*^ the increased level following DA depletion computed with the factor *α* (*α ≥* 1).

The STN D1-like receptors are of D5 sub-type, expressed at post-synaptic sites, and with constitutive activity (Chetrit et al., 2013). They have thus been modeled as modulators of the transfer function of the STN neuron population (see equation next), rather than as modulators of incoming activity:

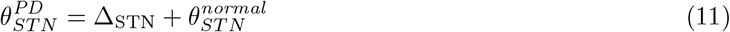

where *θ*_*ST N*_ is the average firing threshold of STN neurons, and Δ_STN_ is the offset created by DA depletion on D5 receptors (Δ_STN_ *≥* 0).

Finally, based on the lack of projective selectivity of D1 and D2 MSN in macaque monkeys, we assumed that, on average, they compensate each other and that, consequently, their influence for the emergence of *β*-band oscillations is non-essential. This simplification constitutes a relatively radical modeling choice, that aims at studying the extent to which PD oscillatory phenomenon can be explained without segregated striatal pathways.

## Code accessibility

The code used to simulate the neural network is available at https://github.com/SN1885A/BCBG-model.

## Acknowledgements

The authors are grateful for the constructive comments of Mark Humphries in an earlier version of this manuscript. This project was partially funded by the ANR EvoNeuro project, ANR-09-EMER-005-01, as well as by the Laboratory of Excellence SMART (ANR-11-LABX-65), supported by French State funds managed by the ANR within the Investissements d’Avenir programme under reference ANR-11-IDEX-0004-02. IC was funded by the Ville de Paris HABOT Project, and the Marie Sklodowska-Curie Research Grant Scheme, Grant Number IF-656262.

